# Longitudinal genomic surveillance of *Plasmodium falciparum* malaria parasites reveals complex genomic architecture of emerging artemisinin resistance in western Thailand

**DOI:** 10.1101/084897

**Authors:** Gustavo C. Cerqueira, Ian H. Cheeseman, Steve F. Schaffner, Shalini Nair, Marina McDew-White, Aung Pyae Phyo, Elizabeth A. Ashley, Alexandre Melnikov, Peter Rogov, Bruce W. Birren, François Nosten, Timothy J.C. Anderson, Daniel E. Neafsey

## Abstract

**Background:** Artemisinin-based combination therapies are the first line of treatment for *Plasmodium falciparum* infections worldwide, but artemisinin resistance (ART-R) has risen rapidly in in Southeast Asia over the last decade. Mutations in *kelch13* have been associated with artemisinin (ART) resistance in this region. To explore the power of longitudinal genomic surveillance to detect signals in *kelch13* and other loci that contribute to ART or partner drug resistance, we retrospectively sequenced the genomes of 194 *P. falciparum* isolates from five sites in Northwest Thailand, bracketing the era in which there was a rapid increase in ART-R in this region (2001–2014).

**Results:** We evaluated statistical metrics for temporal change in the frequency of individual SNPs, assuming that SNPs associated with resistance should increase frequency over this period. After *Kelch13-*C580Y, the strongest temporal change was seen at a SNP in phosphatidylinositol 4-kinase (PI4K), situated in a pathway recently implicated in the ART-R mechanism. However, other loci exhibit temporal signatures nearly as strong, and warrant further investigation for involvement in ART-R evolution. Through genome–wide association analysis we also identified a variant in a kelch-domain–containing gene on chromosome 10 that may epistatically modulate ART-R.

**Conclusions:** This analysis demonstrates the potential of a longitudinal genomic surveillance approach to detect resistance-associated loci and improve our mechanistic understanding of how resistance develops. Evidence for additional genomic regions outside of the *kelch13* locus associated with ART-R parasites may yield new molecular markers for resistance surveillance and may retard the emergence or spread of ART-R in African parasite populations.

## Introduction

Artemisinin-based combination therapy (ACT) is the first-line treatment for *Plasmodium falciparum* malaria infection in most of the world [1,2]. Resistance to ACT treatment, manifested as delayed clearance of parasites following treatment, was first documented in Cambodia in 2009 [3,4] and has since spread throughout SE Asia [5,6]. Mutations in the BTB/POZ or propeller domain of the *kelch13* gene (PF3D7_1343700) have been associated with artemisinin resistance (ART-R), as evidenced by *in vitro* selection [7] and transfection experiments [8], and are associated with reduced cure rates following ACT [9–11]. Surveys have documented the rapid increase in frequency of *kelch13* mutations in SE Asia over the last ten years, driven by a combination of *de novo* mutation creating new resistance alleles and natural selection favoring the spread of existing alleles [5,12–14]. Though *kelch13* resistance mutations are becoming prevalent in SE Asia, and have been observed at low frequency in Africa, South America, and other parts of the world, to date they have not been observed to spread in any location outside of SE Asia [13,15].

There are several hypotheses to explain the failure of *kelch13*-mediated resistance to spread outside of SE Asia via *de novo* mutations or human migration, as has previously occurred with resistance to chloroquine[16] and pyrimethamine [17]. One is that the selective pressure of ACTs on parasite populations is more intense in SE Asia due to lower disease endemicity, commensurately less acquired immunity to disease, and therefore a greater likelihood that infected individuals will become symptomatic and be treated with drugs. This hypothesis is difficult to exclude, but malaria endemicity is highly variable across sub-Saharan Africa and many other regions, and access to ACTs is high in many regions outside of SE Asia [18], making it unlikely that SE Asian drug selection pressure is unique. An alternative hypothesis, to be explored in more detail in this manuscript, is that *kelch13* mutations induce a fitness cost in parasites lacking an appropriate genetic background. In many pathogens, mutations conferring resistance to drugs also confer deleterious fitness effects that are usually suppressed or mitigated by co-segregating compensatory mutations, a phenomenon well documented in bacteria [19,20], yeast [21], and in *P. falciparum* [22–24].

If background mutations that abrogate a fitness cost of *kelch13* mutations or provide resistance to partner drugs are required for the spread of *kelch13* mutations, SE Asia is a favorable location for these mutations to arise and be selected in association with *kelch13* mutations. Beyond the fact that ACTs have been in use for much longer in SE Asia relative to Africa [25], offering a longer window of time for requisite background mutations to occur and rise in frequency, SE Asian parasites experience a lower rate of sexual outcrossing than parasites in most African populations. This is because *P. falciparum* is an obligately sexual but facultatively outcrossing eukaryotic parasite. Meiosis occurs following the union of parasite gametes in the mosquito midgut, but mosquitoes that feed on humans infected by a single genotype of *P. falciparum* will result in self–fertilization of male and female gametes from the same genotype, rather than outcrossing between unrelated genotypes. In low-transmission settings like SE Asia, a majority of human infections are caused by a single parasite genotype [26], leading to infrequent outcrossing relative to high transmission regions like Africa, where human infections may contain multiple parasite genotypes. In addition, there is reduced competition between parasites in low-transmission settings, because most infected humans and mosquitoes harbor only a single parasite genotype. SE Asia, therefore, may be an ideal setting for *kelch13* mutations to maintain association with a favorable genetic background and spread via natural selection.

There is some existing evidence for the involvement of additional loci in ART-R. Two groups have identified a region of chromosome 14 as being associated with slow parasite clearance [27,28]. Miotto et al [29] have suggested that variants in several loci outside of *kelch13* are associated with ART-R.

We hypothesized that background mutations providing compensatory fitness for ART resistance mutations in *kelch13* or mutations at other loci conferring partner drug resistance should rise in frequency over time with *kelch13* resistance mutations. We performed whole genome sequencing of samples collected between 2001 to 2014 from Northwestern Thailand, a period spanning the emergence and spread of ACT resistance in this region [6], using hybrid selection to enrich parasite DNA in early clinical samples collected as dried blood spots without leukocyte depletion. In conjunction with genotype-phenotype association tests and scans for signals of natural selection, we used longitudinal changes in allele frequency to identify a list of candidate mutations that may provide a suitable background for *kelch13* resistance mutations. These markers give insight into the mechanism of *kelch13*-based ART-R, clarify the genomic architecture of this trait, and suggest other loci in the genome that could be informative targets of future ACT resistance surveillance efforts.

## Results

We sequenced a total of 194 isolates distributed among four time intervals between 2001 and 2014 from five clinics situated along the northwestern border of Thailand (Fig. 1). The median clearance rate half-life of the samples sequenced during each time interval exhibits a sharp increase in this region after 2008 [6], indicating that our collection window spans the emergence of ACT resistance in this region (Fig. 1). Although *kelch13* mutations are strongly associated with slow clearance (Additional file 1: Figure S1), clearance rate half-life ranges from 3.0 to 9.6 hours for parasites with *kelch13* mutations, and 16 out of 68 samples with *kelch13* mutations exhibit a clearance rate half-life less than 5 hours, suggesting that resistance may vary according to the nature of different *kelch13* mutations and/or parasite genomic background.

**Fig. 1.**
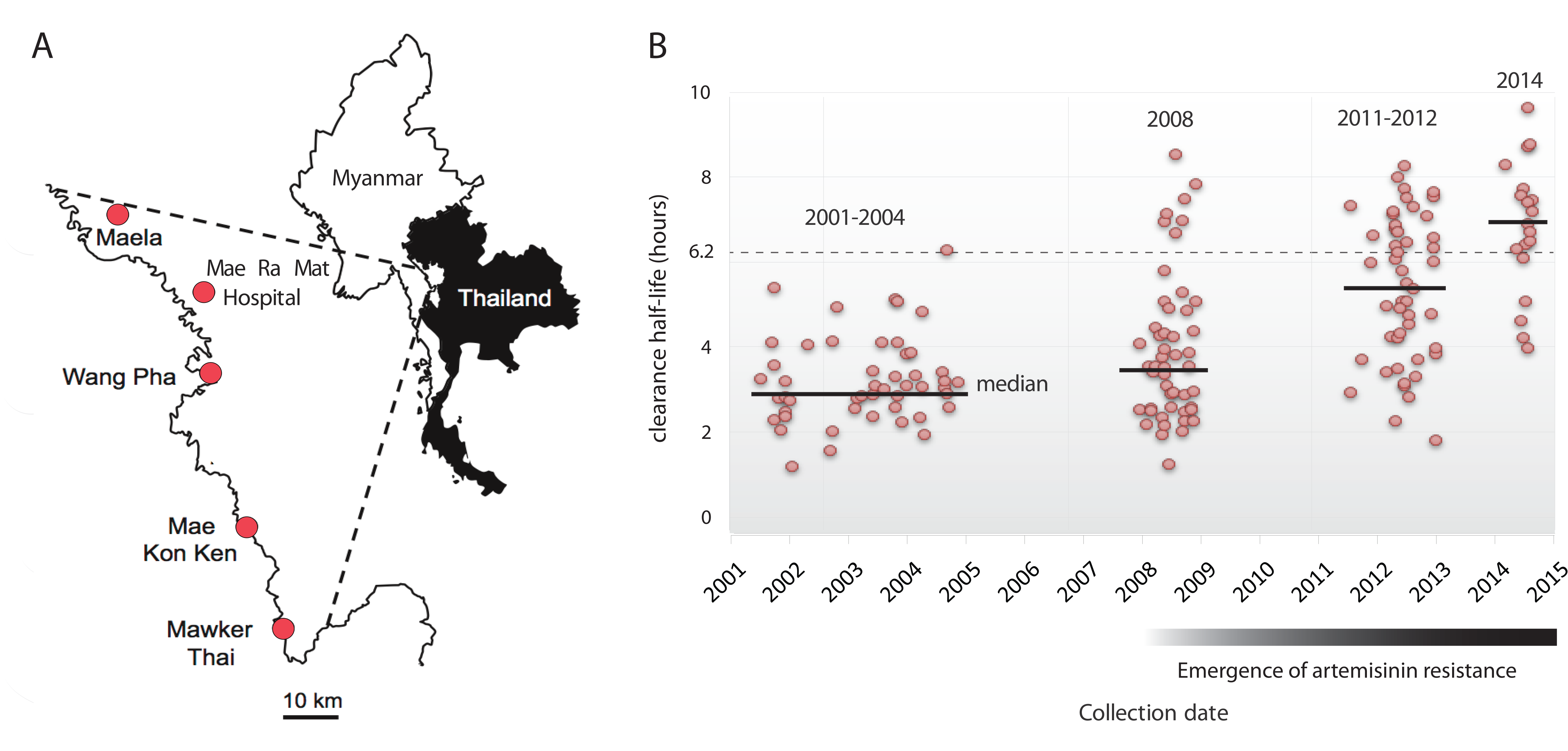
Location, collection date, and parasite clearance rate associated with patient blood samples. A) Location of clinics in the Northwestern Thailand involved in the collection of samples used in this study. B) Distribution of the parasite clearance rate half-life associated with each sample. Horizontal black lines indicate the median clearance rate half-life for each sampling interval.

### Dataset filtration

We performed an analysis of identity by descent (IBD) among pairwise comparisons of samples within sampling intervals to ascertain changes in the degree of clonality due to recent common ancestry over time, using a hidden Markov Model-based approach that makes use of SNP calls [30]. High levels of IBD among samples impedes the identification of individual variants subject to selection due to increased LD among variants within IBD blocks. Analysis of pairwise IBD distributions within each time interval showed only a modest amount of recent common ancestry during the first three sampling intervals, but a high level of clonality among isolates collected in 2014 (Fig. 2). This phenomenon resembles the previously documented increase in parasite clonality in Cambodia, [31], and most likely stems from decreasing disease transmission in this region [26] coupled with the selective sweeps of multiple *kelch13* resistance mutations. The increased IBD among the 2014 samples elevates the LD between SNPs, making it difficult to identify signals associated with individual background mutations. Therefore, the 2014 samples were excluded from subsequent analyses of temporal frequency trends. Other isolates were discarded due to low sequencing coverage depth (see Methods), ultimately resulting in a collection of 134 sequenced isolates for the temporal analysis (Additional file 7: Table S1).

**Fig. 2.**
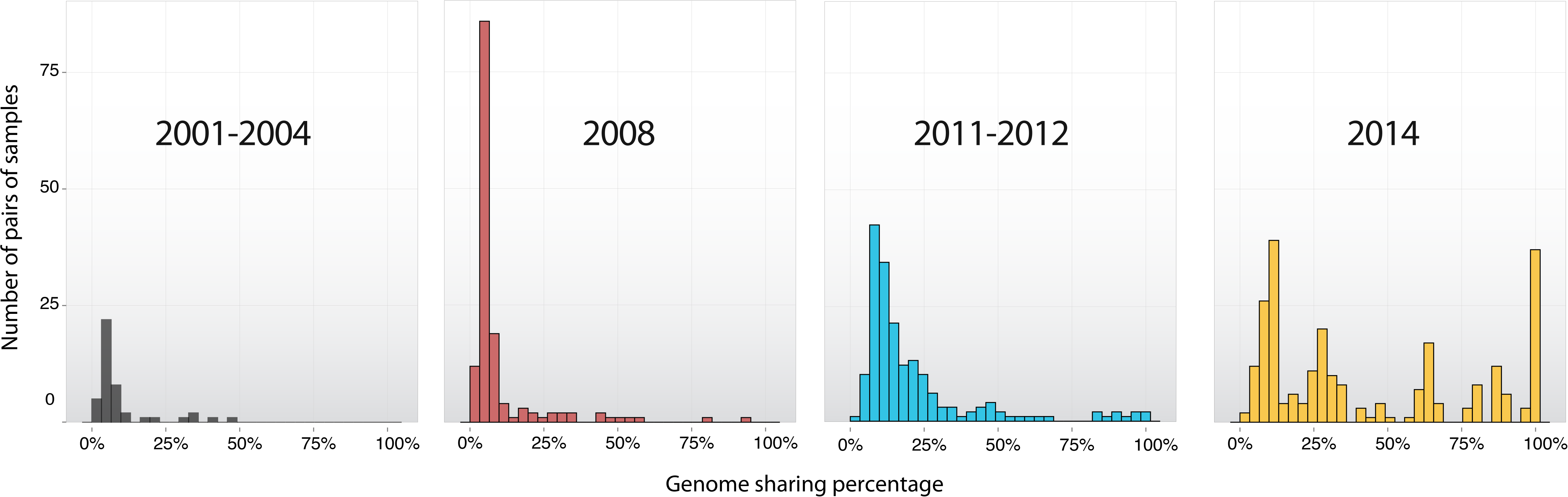
Histogram of the percentage of genome sequence shared between pairs of samples. The vertical axis represents the number of pairwise comparisons exhibiting IBD levels (expressed as a percentage of the full genome) within the range intervals specified on the horizontal axis. Only pairs with a non-zero percentage of genomic sharing are shown.

The dataset was also filtered based on location and nature of polymorphic sites (see Methods). The resultant 15,117 SNPs were analyzed using both a conventional genotype/phenotype association analysis aimed at detecting variants associated with a low parasite clearance rate under artesunate therapy, as well as a phenotype-agnostic approach to identify variants with a temporal trend and other features suggestive of ACT selection. **Frequency trajectory of *kelch13* mutations–** Twelve distinct nonsynonymous *kelch13* mutations were found among sequenced isolates (Additional file 2: Figure S2, Additional file 3: Figure S3A). Figure 3A shows the frequency of *kelch13* mutations exhibiting at least 5% allele frequency in at least one of the first three sampled time intervals. Although other *kelch13* mutations exhibit a higher frequency before 2011, C580Y overtakes those in the 2011-2012 sample. Two independent origins of the C580Y mutation were inferred by the observation of shared haplotypes in the vicinity of that mutation (Additional file 4: Figure S4), consistent with previous SNP genotyping analyses of parasites from the Thai-Myanmar border [32] and other SE Asian locations [12].

### GWAS results

A GWAS using the filtered set of SNPs identified during the first three sampling intervals identified the SNP underlying the *kelch13* C580Y mutation as the variant most significantly associated with slow parasite clearance (P = 7.0e-06; Q = 0.09, Benjamini-Hochberg) [33]. There were 15 other candidate SNPs that, in spite of having q–values greater than 0.10, had p-values lower than expected assuming a uniform distribution of p-values (P < 1e-3; Additional file 5: Figure S5A). This list of SNPs will subsequently be referred to as the GWAS set. Of this set, 11 are nonsynonymous coding mutations and four are synonymous coding mutations (Additional file 7: Table S2). The coding SNPs are located in 13 distinct genes, including three conserved proteins with unknown function and three conserved membrane proteins with unknown function. Three genes containing candidate SNPs encode proteins involved in protein degradation via the proteasome complex, a pathway hypothesized to be associated with ART resistance [34,35]: putative sentrin-specific protease (SENP2) (PF3D7_0801700); ApiAP2 transcription factor (PF3D7_1342900); and a putative ubiquitin protein ligase (PF3D7_1448400). The derived nonsynonymous mutation in the sentrin-specific protease is negatively associated with artemisinin resistance (negative beta value on Additional file 7: Table S2), meaning that the reference allele is associated with slow clearance. For the mutations in the ApiAP2 transcription factor and ubiquitin protein ligase, the derived alleles are positively associated with slow clearance.

An additional association analysis was performed with only those samples containing *kelch13* mutations in a search for other variants that could be potentiating slow clearance rates mediated by *kelch13* mutations. No variants exhibited a statistically significant association with clearance rate after correction for multiple testing. However, the most significant (P = 7.1e-4) non-synonymous SNP positively associated with artemisinin resistance (positive beta-value on Additional file 7: Table S3) is located on chromosome 10 in a gene functionally annotated as “kelch protein, putative” (PF3D7_1022600; *kelch10*). *kelch10* has limited sequence similarity to *kelch13* (Additional file 3: Figure S3B,C), restricted to a few amino acid positions between one of its instances of “Kelch-type beta-propeller” domain (IPR015915) and the “Galactose oxidase, beta-propeller” domain (IPR015916) of *kelch13* [36]. Both domains were defined based on tertiary structure, imported by InterPro from CATH, a protein structure classification database [37], and they represent beta-propellers with 6 and 7 blades in *kelch10* and *kelch13*, respectively, explaining the lack of amino acid sequence similarity between the loci. The mutation in *kelch10* induces a proline to threonine amino acid substitution at position 623 (P623T) and is located between instances of beta-propeller domains (Fig S7B). P623T exhibits variable impact on parasite clearance rate half-life in the presence of different *kelch13* mutations (Additional file 3: Figure S3D). It significantly increases parasite clearance rate half–life in the presence of the *kelch13* E252Q mutation (Wilcoxon rank sum test, P = 8.7e-3) and mildly affects it in the presence of C580Y or other common *kelch13* mutations (P range = 0.23–1). The *kelch10* mutation does not impact clearance rate on a wildtype *kelch13* background (P = 0.07; Additional file 3: Figure S3D). We used PCR-based Sanger dideoxy sequencing to genotype the *kelch10* mutation in 68 additional samples with clearance rate data and the *kelch13* E252Q mutation from the same geographic region. SNP genotyping data (not shown) indicates that 52 distinct parasite clones are represented within these 68 additional samples, with four clonal groups having the P623T mutation on *kelch10*. Clonal groups containing the E252Q (*kelch13*) and P623T (*kelch10*) mutations exhibit a significantly higher clearance rate relative to those with only E252Q mutation (Wilcoxon rank sum test, P = 3.21e-3; Fig. 4). Given that E252Q is the only common *kelch13* mutation in SE Asia that occurs outside the BTP-POZ and propeller domains of *kelch13*, this association may be evidence of an epistatic relationship with *kelch10* that potentiates the resistance phenotype of E252Q. Further functional work will be necessary to elucidate the relationship between these variants.

**Fig 3.**
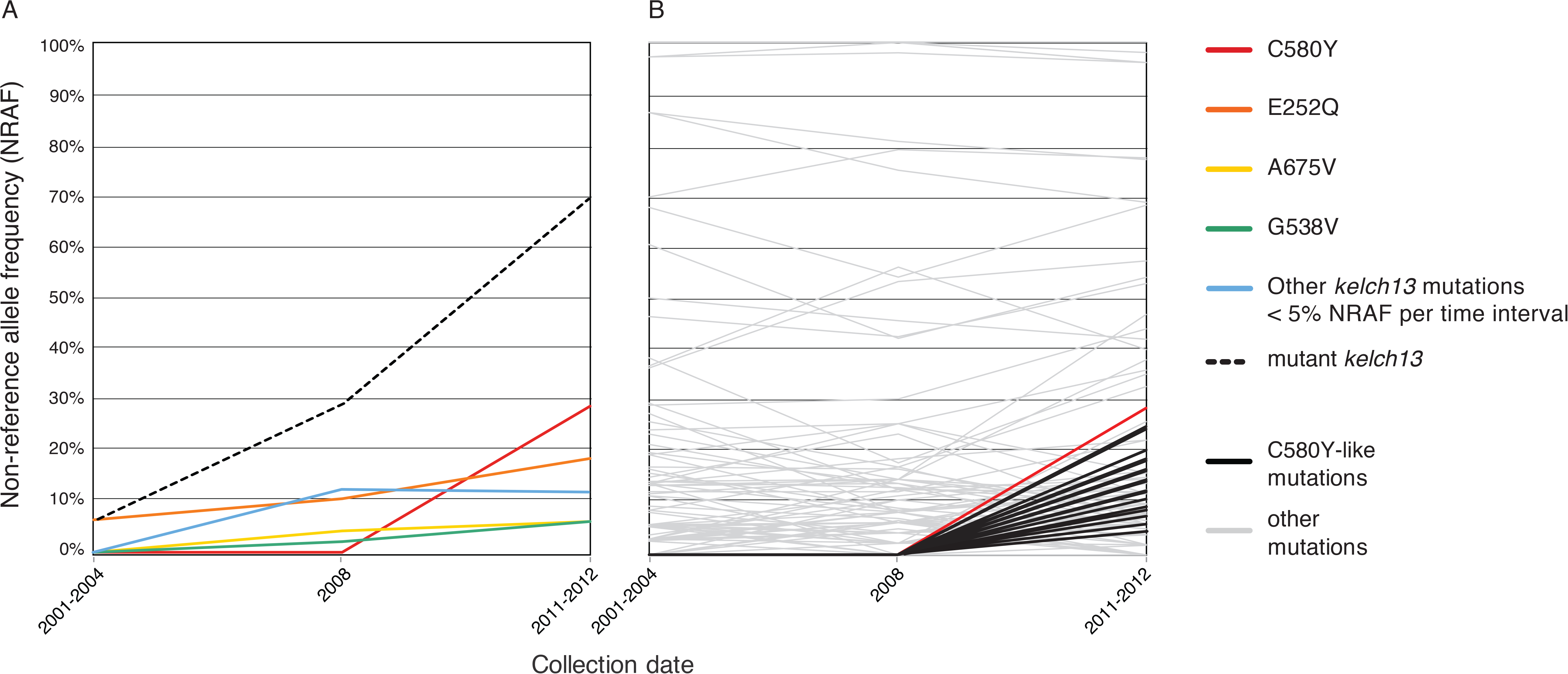
Non-reference allele frequency (NRAF) computed for samples belonging to the first three sampling intervals. (A) NRAF trajectories over time within the *kelch13 resistance* locus. Colored lines indicate the progression of the allele frequency of non-synonymous substitutions located within in kelch13 and having frequency greater than 5% in at least one of the collection eras depicted in this graph (2001-2004, 2008 and 2011-2012) and all other *kelch13* mutation not meeting this criterion (blue line). The dashed line represents the percentage of samples with at least one non-synonymous substitution in *kelch13*. (B) NRAF trajectories over time outside the *kelch13* gene. Gray lines show the frequency of 100 alleles randomly chosen from all single nucleotide polymorphism detected in the dataset. The red line represents the frequency of C580Y across the first three collection intervals. Lines in black indicate the frequency of what we designate as the ‘C580Y-like’ set: alleles absent in the earliest collection phases (2001-2004, 2008), but with NRAF higher than 5% in 2011-2012.

**Fig. 4.**
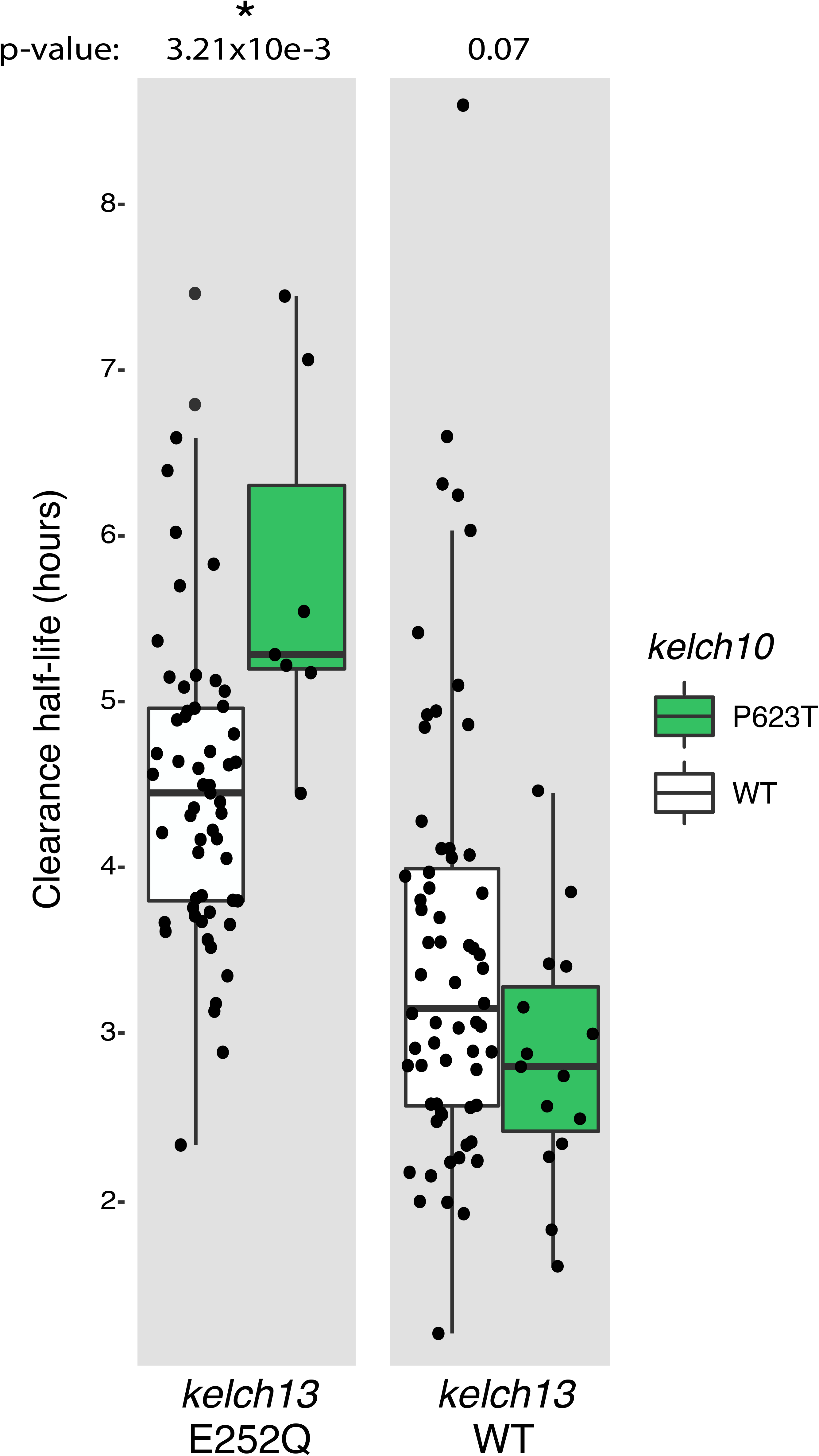
Increased clearance half-life on samples harboring *kelch13* E252Q and *kelch10* P623T mutations. Boxplots on the left illustrate the distribution of clearance half-life on samples with E252Q *kelch13* mutation and either with (green) or without (white) *kelch10* P623T mutation. The boxplot on the right represents the distribution of the same phenotype on samples harboring wild-type (WT) *kelch13*. The difference between distributions shown in each boxplot was evaluated by a Wilcoxon rank sum test.

A paucity of isolates exhibiting wildtype *kelch13* and slow clearance also compromised the power of a GWAS restricted to wildtype *kelch13* isolates. No statistically significant associations were observed after correction for multiple testing.

### Temporal screen for *kelch*13 background SNPs

The set of 15,117 genome–wide SNPs meeting the quality and frequency filters described in the Methods section was screened for positions exhibiting a NRAF trajectory similar to C580Y, under the hypothesis that any variants contributing to a fit genomic background for *kelch13* mutations should increase in frequency in tandem with the most successful *kelch13* mutation. Two approaches were utilized to identify variants with NRAF trajectories similar to C580Y.

The stricter approach, requiring an NRAF of 0% in the first two sampling intervals and an NRAF ≥ 5% in the third interval, yielded 779 SNPs, a set that we designate C580Y-like. An alternative approach, utilizing a three-dimensional vector-distance metric not involving hard NRAF thresholds, yielded 382 SNPs, 158 of which also belong to the C580Y-like set. After removing shared SNPs we designated this set as C580Y-vector-like. Variants segregating non-independently as part of linked haplotype blocks within these two variant sets were identified as described in Methods, and single ‘tag SNPs’ were selected to represent blocks of variants that routinely co-occurred within samples. This reduced the number of variants within the C580Y-like and C580Y-vector-like sets to 254 and 208 independently segregating variants.

Variants in these two lists were next filtered for their degree of association with parasite genotypes harboring mutant *kelch13* genotypes, under the assumption that background mutations under selection for positive fitness effects in conjunction with *kelch13* mutations should be found disproportionately in association with such mutations. Fig. 5A and Additional file 6: Figure S6C illustrate the degree of association of C580Y-like and C580Y-vector-like variants with mutant *kelch13*, relative to panels of control SNPs exhibiting similar NRAF in the third (2011-2012) sampling interval. The candidate background SNPs were binned into quartiles according to their 2011-2012 NRAF. In the C580Y-like set, the lowest quartile of candidate background variants (5% < NRAF ≤ 6%) exhibits no significant difference from control SNPs with regard to LD with mutant *kelch13* (P = 0.47; Wilcoxon rank sum test). The higher frequency quartiles all exhibit enriched LD with mutant *kelch13* relative to frequency-matched control SNPs, however, suggesting they may contain legitimate background variants for resistance mutations at that locus (6% > NRAF ≤ 8%; P = 3.49E-3; 8% > NRAF ≤ 10%, P = 1.14E-3; 10% < NRAF ≤28%, P = 2.23E-8). LD analysis of the C580Y-vector-like set yields qualitatively similar results (Fig S6C).

**Fig 5.**
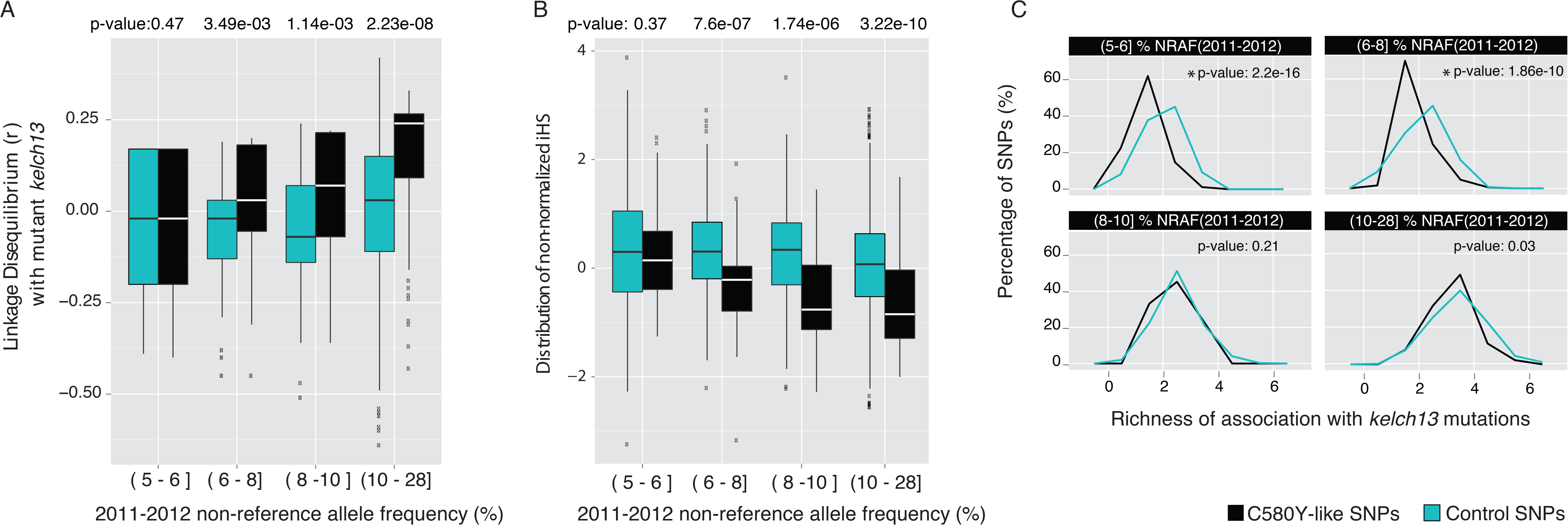
Signatures of selection of C580Y-like SNPs. (A) Distribution of linkage disequilibrium (*r*) between mutant *kelch13* and other SNPs, binned according to the 2011-2012 NRAF (interval left-closed, right-open). Boxplots in black depict the distribution of C580Y-like SNPs. The boxplot in blue depicts the distribution of controls (SNPs with comparable NRAF in 2011-2012, but exhibiting non-zero NRAF in the earlier collection phases). P-values indicate significantly different distributions of C580Y-like and control SNPs (Wilcoxon test). (B) Comparison of the distribution of non-normalized iHS values for C580Y-like SNPs and control SNPs. Lower iHS values indicate a stronger signature of selection. P-values indicate significantly different iHS distributions for C580Y-like vs. control SNPs (Wilcoxon test). (C) Comparison of the *kelch13* allelic richness of association between C580Y-like SNPs (black) and control SNPs (blue). Each graph shows the distribution of richness for SNPs with NRAF (2011-2012) within the same quartiles defined for iHS and LD boxplots; quartiles are indicated on the top of each graph. Horizontal axes indicate the number of distinct *kelch13* mutations co-occurring with each SNP (richness). Vertical axes indicate the percentage of SNPs. Controls consist of SNPs with 2011-2012 NRAF within the same quartile same as candidate SNPs. P-values on the top of the graph indicate significantly different distributions between C580Y-like and control SNPs (Wilcoxon test). The C580Y-like SNPs with NRAF less than 8% generally exhibit association with fewer *kelch13* alleles than control SNPs, indicating they may be more likely to be products of genetic hitchhiking than C580Y-like SNPs with higher NRAF.

Both the C580Y-like and C580Y-vector-like variant sets were also analyzed for signatures of natural selection using the iHS statistic [38], assuming that variants exhibiting upward NRAF trajectories as a result of neutral genetic drift or sampling error would be unlikely to exhibit independent evidence of selection. Similar to the mutant *kelch13* analysis, candidate variants were binned into quartiles according to their 2011–2012 NRAF and compared to control SNPs exhibiting similar NRAF in that sampling interval. Fig. 5B and Additional file 6: Figure S6D illustrate the results of this analysis for the C580Y-like and C580Y-vector-like sets, respectively. As with the analysis for LD with mutant *kelch13,* in the C580Y-like set variants in the lowest 2011-2012 NRAF quartile (5% < NRAF ≤ 6%) exhibited no significant difference in iHS distribution relative to frequency-matched control SNPs (P = 0.37; Wilcoxon rank sum). Once more, however, C580Y-like variants in the three higher NRAF quartiles exhibited significantly lower iHS distributions than control SNPs, suggesting they may be subject to recent natural selection (6% > NRAF ≤ 8%; P = 7.6E-7; 8% > NRAF ≤ 10%, P=1.74E-6; 10% > NRAF ≤28%, P = 3.22E-10; Wilcoxon rank sum test). iHS analysis of the C580Y-vector-like set for natural selection again yields qualitatively similar results (Additional file 6: Figure S6D).

The observation of enhanced LD with mutant *kelch13*, as well as evidence of enriched signals of natural selection in candidate SNPs exhibiting high NRAF in 2011-2012, indicates that some of these variants may constitute part of a genomic background required for parasite fitness in the presence of *kelch13* mutations. An alternative hypothesis is that high-NRAF candidate background SNPs could exhibit these LD and selection signatures as a result of selective sweeps targeting the *kelch13* resistance mutations, and that some or all of these candidate background variants are themselves selectively neutral. Though these candidate markers reside on distinct chromosomes (Additional file 7: Table S4 and S5), reduced opportunities for sexual outcrossing in a low-transmission setting like Thailand could result in selective sweeps that impact the whole genome, rather than just the immediate vicinity of the selected *kelch13* variants on chromosome 13.

To explore this hypothesis, candidate SNPs were examined for the richness of their association with different *kelch13* mutations, under the hypothesis that if these mutations are neutral and were dragged up in frequency due to chance co–occurrence with a *kelch13* mutation undergoing a selective sweep, they should be primarily associated with a single *kelch13* mutation rather than co-occur with multiple different *kelch13* mutations. This analysis indicates that many of the candidate background SNPs with low NRAF exhibit low allelic richness in their *kelch13* mutation association, but that candidate background SNP exhibiting medium to high NRAF (> 8%) are found in association with multiple different *kelch13* mutations, similar to a set of frequency-matched control variants (Fig. 5C), reducing the likelihood that their NRAF increase is due to a trivial *kelch13* hitchhiking scenario. Additional file 7: Table S4 lists all nonsynonymous candidates in the C580Y-like set with NRAF greater than 8%. Additional file 7: Table S5 list nonsynonymous candidates in the C580Y-vector-like set that are disjunct from the C580Y-like set. The Shannon entropy index of the association with distinct *kelch13* mutations was calculated for each candidate. SNPs with a higher Shannon entropy index are less likely to have been detected due to hitchhiking with a single *kelch13* mutation, given their association with multiple distinct *kelch13* mutations (Additional file 7: Table S4 and S5). Table 1 contains a list of some of the most promising candidate background SNPs that pass these filters in the C580Y-like C580Y-vector-like sets, and it includes selected candidates from the GWAS sets. Ranked either by 2011-2012 NRAF or Shannon index, the top candidate in the C580Y-like set is a nonsynonymous mutation in a putative phosphatidylinositol 4-kinase (PI4K) locus (PF3D7_0419900). A distinct *Plasmodium* PI4K beta locus has been identified as a potential *Plasmodium* drug target [39,40], and another component of the phosphatidylinositol pathway (PI3P) has been implicated in the mechanism of *kelch13*–mediated artemisinin resistance [34]. Further functional work will be required to evaluate the potential compensatory role of this and other candidate background mutations, including nonsynonymous mutations in a gene encoding a Sec14-domain-containing protein (PF3D7_0626400), a member of a family of proteins also associated with vesicle trafficking, previously reported as having orthologs in yeast functioning as regulators of PI4K [41] and ranked among the top 14 candidates in C580Y-like set when ranking by Shannon index and 2011-2012 NRAF. Several other genes encoding proteins associated with vesicle trafficking and/or the endoplasmic reticulum were also in the list (Table 1).

**Table 1.**
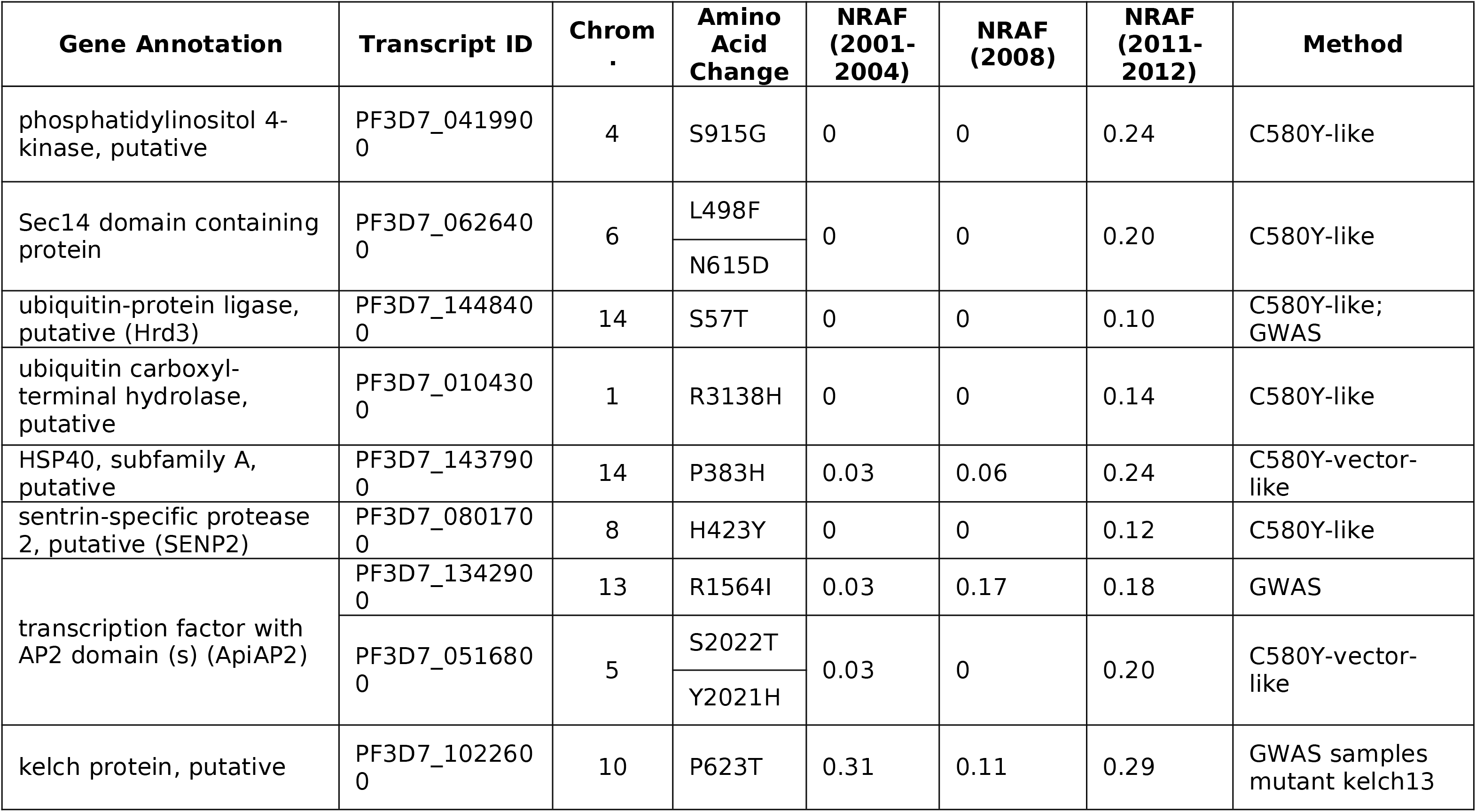
Selection of non-synonymous mutations detected by C580Y-like, C580-vector-like method and SNPs associated with lower clearance rate according to GWAS.

## Discussion

Assuming a conventional eukaryotic mutation rate of approximately 3 × 10-9 mutations per base pair generation [42,43], and assuming that a typical *P. falciparum* infection results in approximately 1011 parasites during its peak, [44] it is likely that virtually all of the 23 million nucleotide positions in the parasite genome mutate into all possible alternative states during the course of a single infection within an individual. The observation that drug resistance does not routinely emerge in populations after years of use of a drug as first line therapy suggests that the genomic architecture of drug resistance that induces minimal fitness cost to the parasite is complex. To be favored by natural selection, resistance mutations must occur on an appropriate genetic background capable of positively potentiating the resistance phenotype and/or compensating for any deleterious fitness impacts that may result from the resistance mutations. Observations of restored susceptibility in parasite populations to anti-malarial drugs within several years of their withdrawal due to high levels of resistance attest to the negative fitness impact of resistance mutations in the absence of drug pressure [45,46].

In this study, we have performed retrospective longitudinal surveillance of the *P. falciparum* genome for mutations inside and outside of the *kelch13* artemisinin resistance locus that may be required for resistance to spread in northwest Thailand. Many of the same longitudinal samples were examined by Cheeseman et al. [47] using a lower resolution targeted SNP genotyping approach in a study that initially identified the chr 13 region adjacent to *kelch13*. Longitudinal genomic surveillance is complementary to GWAS as a means of identifying loci associated with a phenotype subject to natural selection and has the advantage of not requiring phenotype data, which can be much more difficult to collect than genomic data [48–52]. This information potential, combined with the falling cost of genome sequencing, makes a compelling argument for prospective longitudinal genomic surveillance in pathogen populations where the evolution of resistance to therapeutics is a risk.

These results, together with the very detailed longitudinal profile of the temporal dynamics of resistance mutations within the *kelch13* locus itself over the same time interval [14], speak to the complexity of drug resistance evolution when observed in real time instead of after the evolution of a fit resistance genotype. This complexity could give fair warning of the emergence of resistance if changes were observed prospectively via a genome-wide surveillance program. For drugs with resistance profiles requiring multiple mutations, alternation of the drugs used for treatment on a cycle of months or years could disrupt the successful combinations of resistance mutations and genetic backgrounds before either gets too high in frequency in a pathogen population. Drug cycling to deter resistance evolution has been proposed, modeled, and experimentally tested in a bacterial context [53–56], but the effect of the genomic architecture on this kind of therapeutic regimen in a sexually-outcrossing eukaryotic parasite has not been fully explored.

These results also highlight the potential shortcomings of *in vitro* resistance selection studies. While *kelch13* was discovered by sequencing parasites selected *in vitro* for resistance to artemisinin [7], such studies cannot recapitulate the complex fitness landscapes influencing the evolution of parasites exposed to drugs in the field. Mutations observed when sequencing culture-adapted parasite lines selected for resistance to an anti-malarial compound may or may not represent changes that are eligible for spread in parasite populations in the field, because cultured parasites are not exposed to the same fitness-determining challenges as wild parasites. And different wild populations of parasites may assemble a fit resistance genotype in different ways. Notably, we failed to identify any mutations with our GWAS and temporal analyses in NW Thailand that were previously found to be associated with ART resistance in Cambodia [29]. This lack of replication does not impugn the results from the Cambodian study, because there may legitimately be different determinants of fit genomic backgrounds in the two parasite populations. Ideally, markers for resistance and genomic backgrounds that enable resistance should be identified from parasites collected as close as possible to the geographic setting in which surveillance will take place.

The variants we have identified as candidate background mutations required for the spread of *kelch13* resistance mutations occur in some genes that belong to pathways already hypothesized to contribute to ART resistance in *P. falciparum*. Mbengue and colleagues [34] found elevated levels of phosphatidylinositol-3-phosphate (PI3P) to be associated with ART resistance *in vitro*. Elevated levels of PI3P can result from polyubiquitination of phosphatidylinositol-3-kinase (PI3K). We observed no polymorphisms in PI3K, but our top hit in the C580Y-like set was a nonsynonymous mutation in a PI4K locus (PF3D7_0419900), which may impact PI3P or other relevant members of the phosphoinositol pathway in a manner conducive to evolutionarily fit resistance. Dogovski and colleagues [57] identified the proteasome/ubiquitination pathway as associated with ART resistance via that pathway’s involvement in the cellular stress response. They hypothesize that an enhanced stress response, manifested via lower levels of ubiquitination, delays cell death and confers resistance to ART, and observe enhanced resistance *in vitro* when ART is co-administered with proteasome inhibitors. In our GWAS and temporal analyses we find several loci associated with the proteasome/ubiquitination pathway, including a putative sentrin-specific protease (SENP2; PF3D7_0801700) and a putative ubiquitin protein ligase (PF3D7_1448400). These loci, as well as the kelch domain-containing protein from chromosome 10 (PF3D7_1022600) that may be epistatically associated with the E252Q mutation at the *kelch13* locus, constitute a list of potential genetic surveillance targets in other regions of the world where *kelch13* mutations and/or phenotypic resistance are observed, such as Guyana [58].

## Conclusions

Longitudinal genome-wide surveys of *P. falciparum* parasite populations are a powerful tool for identifying markers of resistance and understanding the nature of resistance evolution. Such surveys should be conducted prospectively in pathogen populations where resistance is anticipated to evolve. Our analysis suggests a complex genomic architecture behind the emergence and spread of ART resistance in NW Thailand. The potentially multi-locus nature of high fitness ART-R genotypes decreases the likelihood that such genotypes will emerge *de novo* in parasite populations in sub-Saharan Africa, most of which have a much higher rate of sexual outcrossing concomitant with higher transmission. However, introduction of high fitness Asian ART-R parasites bearing a suite of genomic adaptations into Sub-Saharan Africa could accelerate the establishment of ART-R there, as occurred with pyrimethamine resistance when the triple mutant *dhfr* haplotype was introduced from Asia [17]. The present results justify the continued investment of resources to contain ART-R to those regions of SE Asia where it has shown the capacity to spread.

## Methods

### Sample collection and resistance phenotyping

Details on sample collection and ART phenotyping have been published previously [6]. Briefly, blood samples were collected from Karen and Burmese patients with a high parasite count (>4% infected red blood cells) but no signs of severe malaria. Collection was performed at five clinics (Maela, Wang Pha, Mae Ra Mat, Mae Kon Khen, and Mawker Thai) located along the Thai-Myanmar border (Fig. 1) between 2001 and 2014. Samples collected prior to 2010 were collected as blood spots on filter paper from finger pricks and underwent hybrid selection to enrich parasite DNA prior to sequencing [59]. Samples collected from 2010 onward were collected as venous blood and depleted of leukocytes to reduce host DNA content, and were therefore sequenced without hybrid selection. Approximately 48 samples from each of four time intervals (2001-2004; 2008; 2010-2011; 2014) were selected for sequencing following SNP genotyping to avoid sequencing multiple isolates exhibiting the same parasite genotype[26]. Following treatment with mefloquine-artesunate combination therapy [6], parasite density was monitored at six hour intervals until patients were slide negative, at which point combination therapy was administered. Parasite clearance data were used to estimate clearance half-life as described previously [14]. Ethical approval for this work was given by the Oxford Tropical Research Ethics Committee (OXTREC 562-15) and the Faculty of Tropical Medicine, Mahidol University (MUTM 2015-019-01).

### Library preparation and sequencing

Illumina sequencing libraries containing a 200 bp insert were constructed and samples were sequenced on an Illumina HiSeq 2500 platform using paired-end 101 bp reads. Samples collected prior to 2010 were enriched for parasite DNA using a previously published hybrid selection protocol [59] and a set of RNA oligonucleotide baits prepared by random transcription of genomic DNA prepared from the 3D7 parasite line.

### Genotype calling

Reads were aligned using BWA [60] (sample, default parameters) against the *P. falciparum* 3D7 v3 reference genome assembly (PlasmoDB release 12.0). Fourteen samples exhibiting less than five-fold read coverage depth across 60% or more of the reference assembly were excluded from analysis. Genotype calling was performed using the GATK UnifiedGenotyper (version 2.4-9) [61] using the following parameters: EMIT_ALL_SITES,–stand_emit_conf 0.0,–stand_call_conf 0.0,–sample_ploidy 2 and–glm SNP (SNPs only, no INDELs). Annotation of polymorphic sites and evaluation of their effect on the coding sequence of genes was done via SNPEff [62]

### Genotype filtering

Polymorphic sites meeting any of the following criteria were removed from the analysis using VCFTools [63] and custom scripts: heterozygous sites, QUAL < 60, GQ < 30, indels, sites lacking genotype calls in 10% or more of the samples within one or more time intervals and sites with non-reference allele frequency (NRAF) lower than 5% in all time intervals. NRAF was calculated using custom scripts. Polymorphic sites located in pericentromeric, subtelomeric and hypervariable regions (listed in Additional file 7: Table S6) were removed, as were those occurring in genes belonging to large antigenic gene families (Additional file 7: Table S7).

### Identifying regions identical by descent (IBD)

A previously described Hidden Markov Model (HMM) was utilized to identify Identical by Descent (IBD) genomic regions between pairwise comparisons of samples [30].

### Genome wide association scan (GWAS)

Genotype calls were converted from VCF format to PLINK format using VCFTools [63]. GEMMA [64] was run with default parameters to evaluate the degree of association between genotype calls and the clearance rate half-life under artemisinin treatment. Q-Q and Manhattan plots were rendered using the qqman R package [65]. P–values were corrected for multiple testing using the Benjamini-Hochberg method [33] implemented in the standard R package [66].

### Comparison between *kelch13* and *kelch10*

Protein sequences of the *kelch13* and *kelch10* (PF3D7_1022600) genes were aligned using NCBI-BLAST web page [67]. Domains were identified using InterProScan [36,68]. Projections connecting regions of similarity between the linear representation of *kelch13* and *kelch10* sequences (Additional file 1: Figure S1) were rendered by Kablammo [69]. Boxplots in Additional file 1: Figure S1, as others boxplots in this work, were rendered using ggplot2 R package [70].

### Temporal Analysis

Two methods were employed to identify SNP sets with temporal frequency signatures concordant with selection for ACT resistance. First, SNPs were identified that exhibited a sample frequency history strictly similar to the C580Y *kelch13* mutation (‘C580Y-like’), which has prevailed as the most successful ART resistance mutation in the region [14]. To be defined as C580Y-like, SNPs were required to exhibit a NRAF of 0 during the first two sampling intervals (2001-2004 and 2008) and NRAF ≥ 5% during the third sampling interval (2011-2012).

The second approach to identify SNPs with population histories similar to that of C580Y was less strict with regard to NRAF requirements. SNP frequency changes over time were interpreted as three–dimensional vectors, with each dimension corresponding to the NRAF of the SNP in one of the first three sampling intervals (2001-2004; 2008; 2011-2012). The Manhattan distance was calculated between the vector representing C580Y and the vectors representing all other SNPs. Distances to the C580Y vector were ranked and a cutoff value was applied according to the first plateau in this series (Additional file 7: Figure S8A). The plateau was empirically defined as a consecutive series of at least 30 C580Y vector distances with a slope equal to 0. SNPs with distances less than the first plateau were designated as ‘C580Y-vector-like’.

### Defining haplotype blocks

Linkage disequilibrium was measured by calculating the Pearson correlation coefficient (*r*) between all pairs of SNPs within the C580Y-like and C580Y-vector-like sets using custom scripts. Pairs of SNPs exhibiting *r* greater than or equal to 0.8 were classified as belonging to the same haplotype block.

### Extended haplotype homozygosity

Genotypes were imputed by Beagle[71] using the recombination map described on [72] and default parameters. Integrated haplotype score (iHS) [38] was calculated via the REHH R package[73] using scan_hh command (limhaplo=2, limehh=0.05, limehhs=0.05) and *Plasmodium reichenowi* genome as the ancestral genotype. When ancestral state could not be determined from *P. reichenowi*, the *P. falciparum* 3D7 v3 (PlasmoDB release 12.0) assembly was assumed to reflect the ancestral allele. iHS of triallelic sites (distinct ancestral, derived and reference alleles) were not computed.

### Linkage disequilibrium with mutant *kelch13*

‘Mutant *kelch13*’ was defined as any *kelch13* genotype containing one or more non-synonymous differences relative to the 3D7 reference sequence (Additional file 9: Figure S9A). Linkage disequilibrium (*r*) was calculated between every candidate SNP in the C580Y-like and C580Y-vector-like variant sets and mutant *kelch13*.

### Correlation between 2011-2012 allele frequency with signatures of selection

The median of iHS and the median LD with mutant *kelch13* were calculated for each haplotype block and for each singleton SNP in the C580Y–like and C580Y-vector-like variant sets. Haplotype blocks and singleton SNPs were divided into quartiles according to the distribution of their frequency (median frequency, in the case of haplotype blocks) in the third time interval (2011-2012). The distributions of iHS and LD with mutant *kelch13* for each quartile are represented in Figure 4. Difference between distributions were evaluated by Wilcoxon rank sum test using the standard R package [66]. The haplotype block containing the C580Y mutation was not included in this analysis. Control SNPs were selected for comparison to the C580Y-like and C580Y-vector-like SNP sets according to their frequency in the third time interval (2011-2012), with no requirements regarding observed frequencies in the previous two time intervals.

### Richness of co-occurrence and LD with distinct *kelch13* mutations

Richness was defined as the number of distinct *kelch13* mutations co-occurring with a candidate SNP. The two independent C580Y mutations were considered individual events for this analysis. SNPs with 2011-2012 NRAF fitting each quartile range were used as controls. The distribution of the number of samples exhibiting control SNPs was matched to the equivalent distribution for candidate SNPs within each quartile. Differences between distributions were evaluated by a Wilcoxon rank sum test implemented in the standard R package [66].

To further explore the diversity of *kelch13* allelic association, linkage disequilibrium (*r*) was calculated between each SNP in the C580Y-like and C580Y-vector-like sets and each *kelch13* mutation. Only samples harboring either the *kelch13* mutation being evaluated or wild type *kelch13* were considered on each compute because multiple *kelch13* mutations never co-occur in the same parasite in our dataset. SNPs exhibiting significant LD with at least two *kelch13* alleles was used as a threshold for identification of a set of high-confidence candidates from the C580Y-like and C580Y-vector-like variant sets.

Shannon entropy index was calculated for each candidate according to the number of samples harboring that SNP and each individual *kelch13* mutations. The two independent C580Y mutations were evaluated individually in this analysis.

## List of Abbreviations

ART: artemisinin
ACT: artemisinin combination therapy
SNP: single nucleotide polymorphism
GWAS: genome-wide association study
NRAF: non-reference allele frequency
IBD: identical-by-descent

## Declarations

### Ethics Approval and Consent to Participate

Ethical approval for this work was given by the Oxford Tropical Research Ethics Committee (OXTREC 562-15) and the Faculty of Tropical Medicine, Mahidol University (MUTM 2015-019-01).

### Consent for Publication

Not applicable

### Availability of Data and Material

The dataset supporting the conclusions of this article has been submitted to EuPathDB (http://eupathdb.org) is available via the NCBI BioProject database, accession PRJNA262567.

### Authors’ Contributions

GCC performed the longitudinal, GWAS, and selection analyses and wrote the first draft of the manuscript. DEN, TJCA, and IHC conceived the study, oversaw analysis, and helped to write the manuscript. SFS performed the IBD analysis. FN oversaw sample collection and provided project guidance. APP and EAA collected and processed the clinical samples. BWB oversaw data generation and analysis and provided project guidance. AM and PR performed hybrid selection. SM and MM-W extracted DNA and performed sample genotyping. All authors reviewed and contributed to the writing of the manuscript.

## Acknowledgements

We acknowledge Sinéad Chapman, James Bochicchio, and Caroline Cusick for project management at the Broad Institute. We acknowledge Daniel Park, Allison D. Griggs and Eli Moss for providing computer scripts for read alignment, genotype calling and allele frequency estimation. We also acknowledge Daniel Park for sharing the recombination map used in the genotype imputation for the iHS analysis. We thank the staff of SMRU and the malaria patients on the Thai-Myanmar border for providing the samples used in this study.

## Additional Files

Additional file 1: Figure S1 **Distribution of clearance rate half-life of samples exhibiting wildtype and mutant *kelch13*.**

Additional file 2: Figure S2 **Non-reference allele frequency (NRAF) for** *kelch13* polymorphisms during the first three sampling intervals. (A) Colored vertical bars represent the NRAF of each *kelch13* mutation detected in our dataset and listed on panel B. Diagonal lines connect the boundaries of the bars representing the same mutation across the three sampling intervals. (B) List of the NRAF of each *kelch13* mutation and the NRAF of “mutant *kelch13*” (sum of the NRAF of all *kelch13* mutations) observed during the first three sampling intervals.

Additional file 3: Figure S3 **Location of** ***kelch13*** **and** ***kelch10*** **mutations and clearance half-life of samples harboring those mutations**. (A) Location of SNPs on *kelch13* gene (horizontal axis) and clearance half-life of each sample collected in the first three time intervals and harboring those mutations (vertical axis). (B) Location of InterPro domains identified on *kelch13* and *kelch10* amino acid sequence. The position of P623T mutation on *kelch10* is indicated by a black diamond. (C) Results of BLAST similarity search between *kelch13* (query) and *kelch10* (subject) amino acid sequence. (D) Box plots showing the distribution of clearance half-life of samples harboring wild-type (WT) and mutant *kelch13* (affected amino-acid listed on the bottom) and either with (green) or without (white) P623T *kelch10* mutation.

Additional file 4: Figure S4 **Evidence of two independent origins C580Y allele.** Each row corresponds to a sample in our dataset harboring a C580Y mutation. Columns represent all polymorphic sites in the vicinity of C580Y allele. Colored circles indicate the genotype according to the legend on the left. Plus signs indicate positions with distinct genotypes between the two haplotypes blocks.

Additional file 5: Figure S5 **Results of association analysis of genotype as predictors of clearance rate.** A) Q-Q plot comparing the expected distribution of p-values and the observed distribution based on the Wald test implemented by GEMMA package [64]. SNPs with p-value lower than10-3 were deemed relevant based on their departure from the diagonal line that extrapolates identity between observed and expected p-values. C580Y has the lowest p-value among all SNPs evaluated. B) Manhattan plot indicating the location of SNPs along the 14 nuclear chromosomes of *P. falciparum*.

Additional file 6: Figure S6 **Selection on C580Y-vector-like candidate SNPs.** (A) Graph indicating the slope of the ranked distance between C580Y vector and others SNPs in the dataset (see Materials and Methods for details). Only the 500 closest SNP-vectors to C580Y-vector are shown. Rectangle in red indicates a contiguous stretch of 30 data points with slope equal to 0. (B) Trajectory of the allele frequency of C580Y-vector-like SNPs across the first three time intervals. (C) Distribution of linkage disequilibrium (*r*) between mutant *kelch13* and C580Y-vector-like (black) and control SNPs (blue), binned according to the 2011-2012 NRAF (interval left-closed, right-open). Controls SNPs have comparable NRAF in 2011-2012, but exhibit non-zero NRAF in the earlier collection phases. P-values indicate significantly different distributions between C580Y-like and control SNPs (Wilcoxon test). (D) Comparison of the distribution of non-normalized iHS values for C580Y-vector-like SNPs and control SNPs. Lower iHS values indicate stronger signature of selection. P-values on the top of the graph indicate significantly different distributions of C580Y-like and control SNPs (Wilcoxon rank sum test).

Additional file 7: Supplementary Tables S1-7. (XLSX 623 Kb)

